# Gastrin producing syngeneic mesenchymal stem cells protect non-obese diabetic mice from type 1 diabetes

**DOI:** 10.1101/2021.05.13.444059

**Authors:** Marie-Claude Gaudreau, Radhika R. Gudi, Gongbo Li, Benjamin M. Johnson, Chenthamarakshan Vasu

## Abstract

Progressive destruction of pancreatic islet β-cells by immune cells is a primary feature of type 1 diabetes (T1D) and therapies that can restore the functional β-cell mass are needed to alleviate disease progression. Here, we report the use of mesenchymal stromal/stem cells (MSCs) for the production and delivery of Gastrin, a peptide-hormone which is produced by intestinal cells and fetal islets and can increase β-Cell mass, to promote protection from T1D. A single injection of syngeneic MSCs that were engineered to express Gastrin (Gastrin-MSCs) caused a significant delay in hyperglycemia in non-obese diabetic (NOD) mice compared to engineered control-MSCs. Similar treatment of early-hyperglycemic mice caused the restoration of euglycemia for a considerable duration, and these therapeutic effects were associated with protection of, and/or higher frequencies of, insulin producing islets and less severe insulitis. While the overall immune cell phenotype was not affected profoundly upon treatment using Gastrin-MSCs or upon in vitro culture, pancreatic lymph node cells from Gastrin-MSC treated mice, upon ex vivo challenge with self-antigen, showed a Th2 and Th17 bias, and diminished the diabetogenic property in NOD-*Rag1* deficient mice suggesting a disease protective immune modulation under Gastrin-MSC treatment associated protection from hyperglycemia. Overall, this study shows the potential of production and delivery of Gastrin in vivo, by MSCs, in protecting insulin producing β-cells and ameliorating the disease progression in T1D.

## Introduction

Progressive destruction of pancreatic islet β-cells by immune cells, which ultimately results in the inability of pancreas to produce sufficient amounts of insulin, is the key event responsible for T1D^[1,2]^. The ideal goals of clinical intervention strategies for T1D are: 1) to prevent or arrest the onset and progression of autoimmunity before hyperglycemia is detected in at-risk individuals and 2) to restore insulin producing β-cell mass and glyco-metabolic and immune homeostasis after clinical diagnosis in T1D patients. Although immuno- and cellular- therapies are widely being tested for their efficacy in preventing/delaying T1D in pre-clinical models and clinical trials[3–10], there is an urgent need for safe and effective approaches to prevent the disease in at-risk subjects and restore insulin independence in T1D patients.

Recently, there has been a major focus on cellular therapy to generate β-cells for T1D treatment[11–13]. In addition, the immune-modulatory properties of some types of stem cells[14–17] can be exploited to suppress autoimmunity. Mesenchymal stromal/stem cells (MSCs), which can be isolated from a number of sources including the bone marrow (BM) and adipose tissue, are adult stem cells with a potential for differentiation into various tissues and they have the capacity for self-renewal and differentiation with a broad tissue distribution, and the ability to repair tissue damage[18–22]. Natural ability of MSCs to migrate in the body has been described[23–25]. Of particular interest for tissue repair, i.v. injection of MSCs results in their specific migration to a site of injury or inflammation[26–29]. Notably, most MSC based approaches for T1D diabetes therapy have been focused on their multi-potency to become insulin-producing cells (IPCs) or immune modulatory properties[30–35]. However, a major caveat is that MSCs from pre-clinical models as well as T1D patients show gross defects in many of these properties and lack the ability to promote protection from T1D[34,36], raising the concern that unaltered autologous MSCs may not have therapeutic value. Studies in NOD mice have shown that repeated injections with allogeneic or xenogeneic MSCs, but not syngeneic MSCs, promote protection from T1D[15,34,37].

One potential solution for overcoming the inherent defects of autologous/syngeneic MSCs is by engineering them to express desired factors that can protect/preserve β-cell function. Here, we examined the potential of syngeneic MSCs that are engineered to express Gastrin, a peptide hormone that is produced by intestinal cells as well as fetal islet hormone and has the ability to stimulate β-cell neogenesis and increase islet mass[38–41], to prevent T1D incidence and reverse the early-stage hyperglycemia in NOD mice. We show that, while non-engineered and control engineered MSCs had no substantial impact on T1D incidence in NOD mice, a single injection of Gastrin-MSCs 1) at pre-diabetic stage resulted in the prevention of hyperglycemia and 2) at early-diabetic stages reversed hyperglycemia, for significant durations. We also show that Gastrin-MSC recipients had profoundly higher frequencies of insulin-producing islets compared to control-MSC recipients, suggesting that the Gastrin-MSC treatment preserves/suppresses the loss of functional β-cell mass and/or induces the production of insulin by residual β-cells. While Gastrin-MSC treatment had no profound impact on global Treg, IL10+ and IFNγ+ T cell frequencies, PnLN cells from these mice showed, as compared to control-MSC treated mice, a significant difference in the cytokine profile, upon challenge with β-cell antigen ex vivo. Overall, this study shows the potential of using autologous MSCs for in vivo production and delivery of growth factors and peptide hormones such as Gastrin in protecting and/or increasing insulin producing β-cell mass to prevent and/or suppress the progression of T1D.

## Materials and methods

### Mice

Wild-type NOD/ShiLtJ (NOD), NOD-BDC2.5-TCR-transgenic, and NOD-*Rag1-/-* mice, originally purchased from the Jackson laboratory (Maine, USA), were bred in our facility. All animal studies were approved by the animal care and use committee of MUSC. To detect hyperglycemia in mice, glucose levels were determined in the blood collected from tail vein, at timely intervals using the Ascensia Micro-fill blood glucose test strips and an Ascensia Contour blood glucose meter (Bayer, USA). Mice with glucose level of >250 mg/dl for two consecutive testing (at weekly time points) were considered diabetic/hyperglycemic. Untreated mice with glucose levels <100 mg/dl (10-12 weeks of age), 140-250 mg/dl, >450 mg/dl were considered pre-diabetic, early-hyperglycemic and overt-hyperglycemic respectively. In studies using mice at various stages of hyperglycemia, “high” glucose reading by Contour glucose meter for 2 consecutive testing were considered the endpoint.

### Peptide antigens and other reagents

Immunodominant β-cell antigen peptides [viz., 1. Insulin B_(9-23)_, 2. GAD65_(206-220)_, 3. GAD65_(524-543)_, 4. IA-2beta_(755-777)_ and 5. IGRP_(123-145)_ were custom synthesized (GenScript Inc) and used in this study. These peptides were pooled at an equal molar ratio and used as β-cell-Ag peptide cocktail as described in our earlier studies[42–46]. PMA, ionomycin, Brefeldin A, ELISA and unlabeled and fluorochrome labeled antibodies, and other key reagents were purchased from Sigma-Aldrich, BD Biosciences, eBioscience, Invitrogen, Millipore, Miltenyi Biotec, StemCell Technologies, R&D Systems, Biolegend, and Santa-Cruz Biotechnology. Magnetic bead based 26-plex cytokine kits were purchased from Invitrogen (Catalog # EPXR260-26088-901). These Luminex multiplex assay plates were read using the FlexMap3D instrument. Gastrin EIA kit was purchased from Sigma-Millipore. Flow cytometry data was acquired using FACS Calibur, FACS Verse or CyAn-ADP instruments and analyzed using Flowjo application.

### Cloning and Lentiviral production

The third-generation replication-incompetent lentiviral cDNA cloning vector (pCDH1-copGFP) and packaging plasmids (supplemental Fig. 1) were purchased from SBI and used as described in our previous report[47]. For cloning Gastrin cDNA, pCDH1-copGFP vector was digested using NheI (downstream of CMV promoter) and NotI restriction enzymes, and the restriction digestion fragment was replaced with PCR amplified and NheI and NotI digested mouse Gastrin cDNA. mRNA isolated from mouse stomach tissue was used to generate the Gastrin cDNA. For control (GFP) and Gastrin lentiviral production, HEK293T packaging cells were transfected with above control and Gastrin-cDNA vectors along with packaging vectors by calcium phosphate method. Occasionally, GPRG cells[48], provided by National Gene Vector Biorepository, were used as virus packaging cells. Virus-containing media were collected after 36 and 72 h, pooled, and centrifuged at 3500 RPM for 15 min and the supernatants were subjected to 0.22-μm filtration, and concentrated at least 25-fold by PEG precipitation. The infectious titers/transduction unit (TU) of the virus were determined by using fresh HEK293T cells.

### Culture, lentiviral transduction, and characterization of bone marrow (BM) MSCs

BM cells were collected from the femurs and tibiae, red blood cells were removed by hypotonic lysis, and washed in complete DMEM/F12 medium containing 10% FBS, antibiotics/antimycotics, L-glutamine, sodium pyruvate, and non-essential amino acids. The cells were seeded in cell culture treated dishes at 2×10^6^ cell/ml. Non-adherent cells were removed after 48h and 72h of cultures, trypsin-treated adherent cells were split every 3^rd^ day starting day 5, and 13-16 day old cultures were used for lentiviral transduction and phenotypic characterization. For transducing the MSCs to obtain >80% efficiency, cells were seeded along with a final concentration of at least 5×10^8^ TU/ml of virus in the presence of polybrene (8 μg/ml) and protamine sulfate (10 μg/ml) for up to 24h, washed and cultured in fresh medium for 48h, split at 1:2 ratio and cultured for an additional 48h. Transduction efficiency and cell surface marker expression profile were determined after 72h of transduction by microscopy and/or flow cytometry. Gastrin secretion was determined in post-transduction supernatants, collected at 72h time-points, by using Gastrin EIA kit from Millipore-Sigma.

### In vitro coculture and antigen presentation assay

Immunological assays were carried out as described in our previous studies[44,47]. LPS exposed bone marrow derived dendritic cells (BM DCs) were cultured in the presence of control- and Gastrin- MSCs for 24h and examined for surface expression levels of activation markers such as CD80, CD86, CD40 and MHC II by FACS. Antigen presentation assays were performed by culturing BDC2.5 peptide-pulsed, LPS exposed BM DCs and unlabeled or CFSE-labeled CD4+ T cells purified from the spleens of NOD-BDC2.5-TCR-Tg mice for up to 4 days. In some assays, control- or Gastrin- MSCs were added to these antigen presentation assay wells to assess the impact of Gastrin on T cell activation. CFSE dilution, as an indication of T cell proliferation, was assessed after 4 days of culture. T cells from these primary cultures were examined for Foxp3+ T cell frequencies by FACS. Supernatants from these antigen presentation assay wells were examined for the levels of cytokines, IL2 and IFNγ by Luminex multiplex assay.

### Treatment of NOD mice with engineered MSCs (eMSCs)

Ten-to-12-week-old pre-diabetic and 12-25-week-old early-hyperglycemic, diabetic, overt-diabetic mice were injected i.v. with non-engineered MSCs, control (GFP)-MSCs or Gastrin-MSCs (approx. 1×10^6^ cells/mouse). Mice from multiple cages were pooled and randomly picked for treatment using control- and Gastrin- MSCs. Treated and untreated control mice were monitored for hyperglycemia by testing for blood glucose levels every week. Cohorts of mice were euthanized at different time-points to determine the degree of Gastrin production in the pancreas, insulitis, immune cell phenotype and auto-antigen specific response.

### Adoptive T cell transfer experiment

Fresh PnLN cells from control- and Gastrin-MSC treated mice were cultured ex vivo for 24h in the presence of anti-CD3 antibody (2 μg/ml) for 24h, and transferred into 6-week-old NOD-*Rag1-/-* mice (i.v.; 1×10^6^ cells/mouse) and tested for blood glucose levels every week.

### Histochemical and immunofluorescence analysis of pancreatic tissues

Pancreatic tissues were fixed in 10% formaldehyde, 5-μm paraffin sections were made, and stained with hematoxylin and eosin (H&E) to assess insulitis. Stained sections were analyzed using a 0-4 insulitis severity grading system as described in our earlier studies[45,49,50]. In some experiments, pancreatic sections were stained using anti-insulin (mouse monoclonal, sc-8033 Santa Cruz Biotechnology) and/or anti-glucagon (rabbit polyclonal; Santa Cruz Biotechnology antibodies, followed by Alexa fluor 488- and 568-linked secondary antibodies and DAPI, and scored for insulitis based on DAPI-positive cells in islet areas and insulin positivity. In some experiments, insulin positive islets among total islet structures in different groups were compared.

### Statistical analysis

Mean, SD, and statistical significance (*p-value*) were calculated using GraphPad Prism, Microsoft Excel, and/or online statistical applications. Unpaired *t*-test or Mann-Whitney test was employed, unless specified, for values from *in vitro* and *ex vivo* assays. Fisher’s exact test was used for comparing the total number of severely infiltrated islets (grades ≥3) relative to total number of islets with low or no infiltration (grades ≤2) in test vs. control groups. Log-rank analysis (http://bioinf.wehi.edu.au/software/russell/logrank/) was performed to compare T1D incidence (hyperglycemia) of the test group with that of respective control group. A *P* value of ≤0.05 was considered statistically significant.

## Results

### Engineering of MSCs to express Gastrin and phenotypic characterization of MSCs

For generating Gastrin-expressing MSCs, tissue culture plate-adherent BM cells from NOD mice were cultured for up to 16-days with splitting at least every 3^rd^ day. Cells with MSC phenotype from 13-16-day old culture were used for various experiments. The cells were transduced with control- or Gastrin- lentivirus particles, both with GFP reporter, **(Supplemental Fig. 1)** and cultured for 72h before use in various assays. Transduction efficiency, indicated by GFP expression, was confirmed by fluorescence microscopy **(Fig. 1A)**. In some experiments, secreted levels of Gastrin in control- and Gastrin-lentivirus transduced/engineered MSC (eMSC) culture wells were examined using EIA kit **(Fig. 1B)**. Examination of cell surface expression of MSC related markers confirmed that control and Gastrin transduced cells maintained their MSC-phenotype after lentiviral transduction **(Fig. 1C)**. These results show that BM MSCs from NOD mice can be engineered effectively to express Gastrin, without affecting their phenotype.

**Figure 1:**
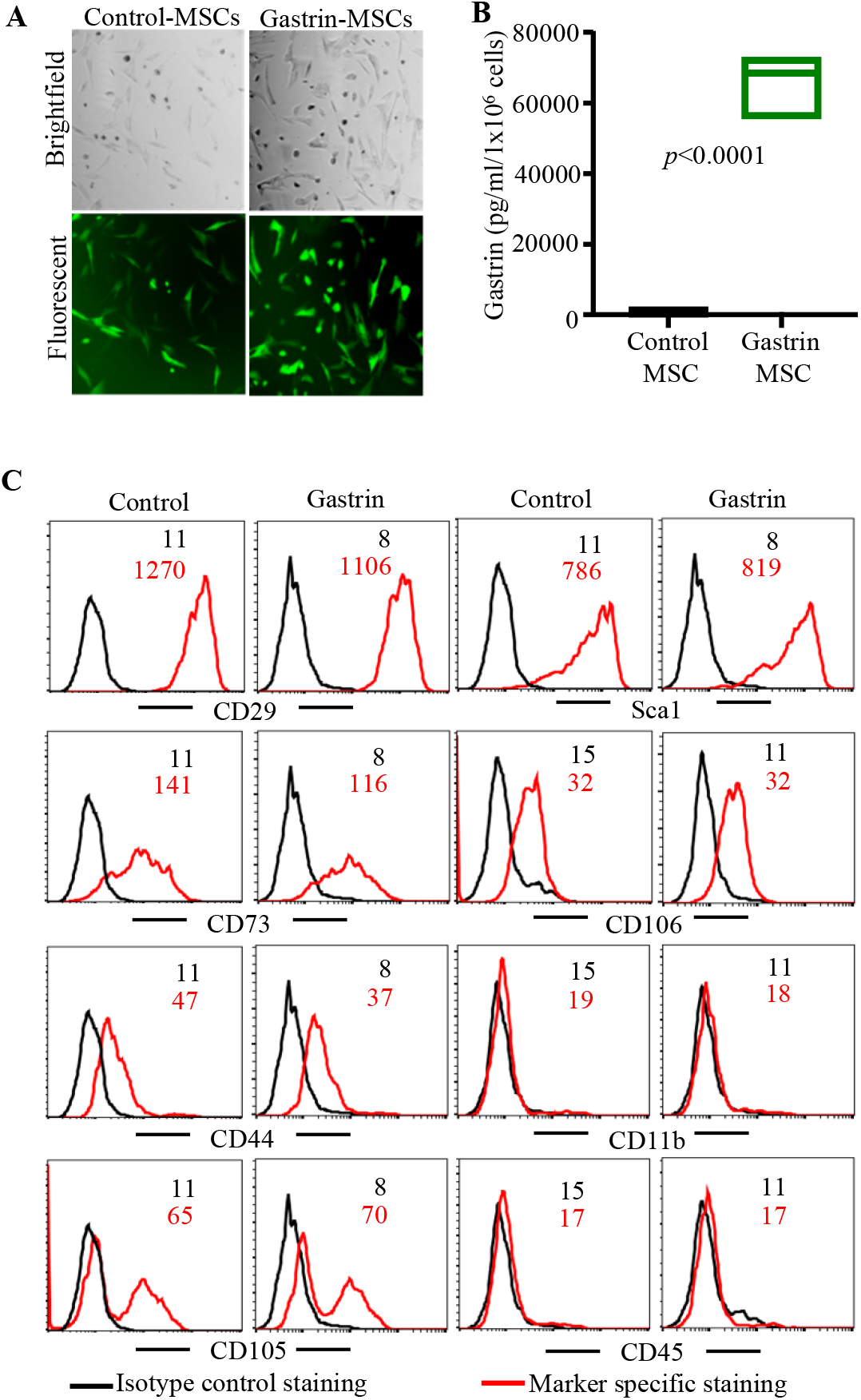
Ectopic expression of Gastrin and phenotypic characterization of eMSCs. BM MSCs from NOD mice were transduced with control vector (GFP) or Gastrin lentiviral particles for 72h as described under materials and methods. **A)** Representative microscopy images of control-MSCs and Gastrin-MSCs are shown. **B)** Supernatants from cells (5×10^5^/ml) cultured for 72h were subjected to competitive EIA and values of supernatants from three independently transduced MSC preparations are shown. *p*-value by paired *t*-test. **C)** eMSCs were stained for MSC markers (CD106, CD105, CD44, CD73, Sca-1 and CD29), and hematopoietic and myeloid cell markers (CD45 and CD11b) and analyzed by FACS. FACS plots and mean fluorescent intensity (MFI) values, representative of three independent assays, are shown.

### eMSCs and Gastrin secretion are detected in pancreatic tissues of treated mice

To assess if the inoculated eMSCs reach pancreas, control-MSCs were injected into pre-diabetic (10-12-weeks-old) or early hyperglycemic (blood glucose: 140-250 mg/dl) NOD mice, 2 or 30 days prior to euthanasia. The frequencies of GFP+ cells in RBC-lysed blood and single cell suspensions of spleen, collagenase and trypsin digested pancreas and kidney were examined. As shown in **Fig. 2A**, considerably higher frequencies of GFP+ cells were detected in the pancreas, compared to blood and spleen, in mice that received eMSCs two days prior to euthanasia. Further, small number GFP+ cells were detectable in the pancreas, even in mice that had received cells 30 days earlier, albeit diminished compared to those which received cells two days before euthanasia. GFP+ cells were not detectable in the kidney, even in mice that received eMSCs two days ahead (not shown). Various tissues including pancreas and kidney were harvested from mice two days post eMSC inoculation, and several intermittent cryosections were examined for green-fluorescence positive cells. While such cells were not detectable in any of the examined kidney sections (not shown), small number of these cells were detected in the pancreatic islets and ductal areas, particularly in those with insulitis **(Fig. 2B)**. To examine if systemically injected Gastrin-MSCs secrete Gastrin in the pancreatic microenvironment and if the Gastrin-secretion persists, early-hyperglycemic NOD mice were injected i.v. with control- and Gastrin-MSCs. Cohorts of mice were euthanized at different time-points, and pancreatic tissues were digested using collagenase and trypsin. Washed single cell suspension of each whole pancreas was cultured in complete DMEM/F12 medium for 48h and the spent media, along with serum samples, were tested for Gastrin levels by EIA. As shown in **Fig. 2C**, profoundly higher levels of Gastrin were detected in the pancreatic suspension cultures of mice that received Gastrin-MSCs compared to those of control-MSC recipients. Interestingly, Gastrin was detectable, albeit at much lower levels, in the pancreas even after 30-days of treatment. These results suggest that eMSCs reach the pancreatic microenvironment, persist, and secrete the factors of interest for a considerable duration. Of note, eMSCs or Gastrin secretion were not detectable in the pancreas of a cohort of mice that were treated at pre-diabetic stage with eMSCs, 90 days prior to euthanasia (not shown).

**Figure 2:**
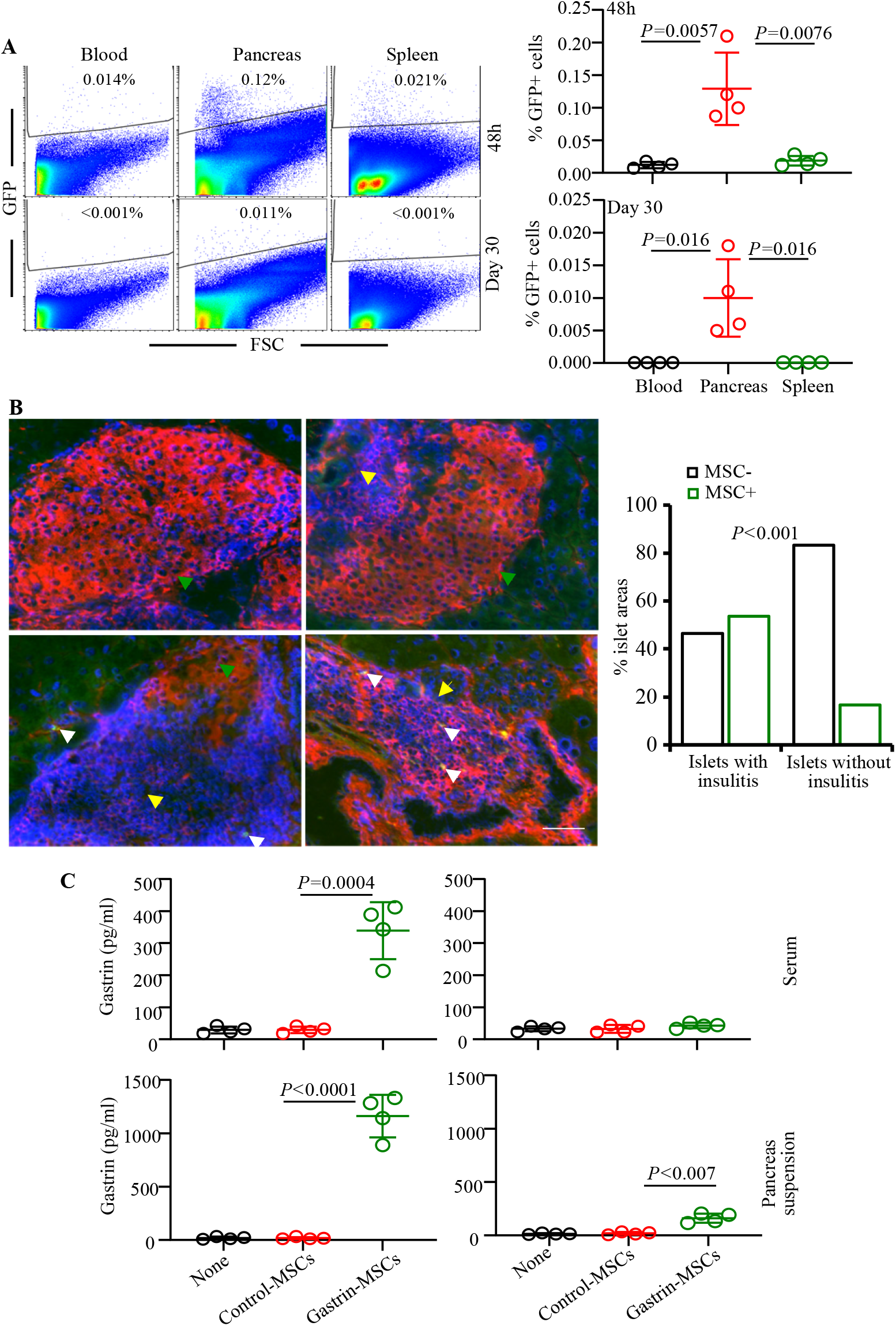
eMSC trafficking, and Gastrin production in the pancreatic tissues of Gastrin-MSC treated mice. Pre-diabetic (10-12 weeks of age) or early-hyperglycemic (12-25 weeks of age; blood glucose: 140-250 mg/dl) female NOD mice were left untreated (none), or injected with control MSCs (1×10^6^ cells/mouse) at different time-points prior to euthanasia. **A)** Blood and various other tissues were processed for flow cytometry to detect GFP+ cells. Representative FACS plots (left panel) of mean±SD values from 4 mice (right panel) that received cells at early-hyperglycemic stage are shown. **B)** Intermittent pancreatic tissue frozen-sections from pre-diabetic and early-hyperglycemic mice, euthanized 48h post control-MSC inoculation, were stained using mouse anti-insulin antibody (β-cells; red) and DAPI (nuclei; blue), and examined for green fluorescence positive cells, and islet and ductal areas were imaged (left panel) using 20x objective. Scale bar: 50μm. Percentage values of islet areas with and without injected MSCs (at least 20 islet areas/pancreas; 3 mice/group) were also shown (right panel). Green and yellow arrows indicate islet areas with strong insulin staining and immune cell infiltration respectively. White arrows point to green fluorescence positive cells. Please note that lower right image shows mostly destroyed islet/ductal area with little or no specific staining for insulin, but with a strong high level of (non-specific) binding by anti-mouse IgG secondary antibody. *P*-value by Fisher’s exact test comparing the presence of GFP+ cells in islets with and without insulitis. **C)** Pancreatic tissues and blood were harvested from cohorts of mice that received control MSCs or Gastrin MSCs 2 or 30 days ahead. Single cell suspensions of whole pancreas were cultured for 24h, and the supernatants along with serum samples were subjected to EIA to determine Gastrin levels. Gastrin values (total amount per pancreas or per ml serum) of 4 mice/group, each tested separately, are shown. *p*-value by Mann-Whitney test.

### Gastrin-MSC treatment at pre-diabetic stage results in diminished insulitis and prevention of T1D in NOD mice

To determine the effect of treatment with Gastrin-MSCs on insulitis and T1D incidence, pre-diabetic female NOD mice were injected once with control-MSCs or Gastrin-MSCs, and the degree of insulitis was determined after 30 days. H&E staining of pancreatic sections revealed that while mice that received control-MSCs did not show significant differences in the insulitis severity compared to untreated controls, mice that received Gastrin-MSCs had significantly reduced insulitis compared to control-MSC recipients (**Fig. 3A**). To assess the impact of suppressed insulitis in Gastrin-MSC recipients on diabetes incidence, cohorts of treated mice were monitored for hyperglycemia for up to 25-weeks post-treatment. As shown in **Fig. 3B**, while 100% of Gastrin-MSC treated mice remained euglycemic for at least 10 weeks post-treatment, more than 60% of untreated and control-MSC treated mice turned hyperglycemic during this period. Moreover, about 40% Gastrin-MSC recipients remained euglycemic at the end of 25 weeks of monitoring. Importantly, diabetes onset was detected in the control groups of mice within 2-5 weeks of monitoring as compared to 12 weeks post-treatment in Gastrin-MSC treated mice. Overall, these results show that a treatment with a single dose of Gastrin-MSCs can result in the retention of significant number of intact islets or islets with less severe insulitis, and prevention of hyperglycemia for a significant duration.

**Figure 3:**
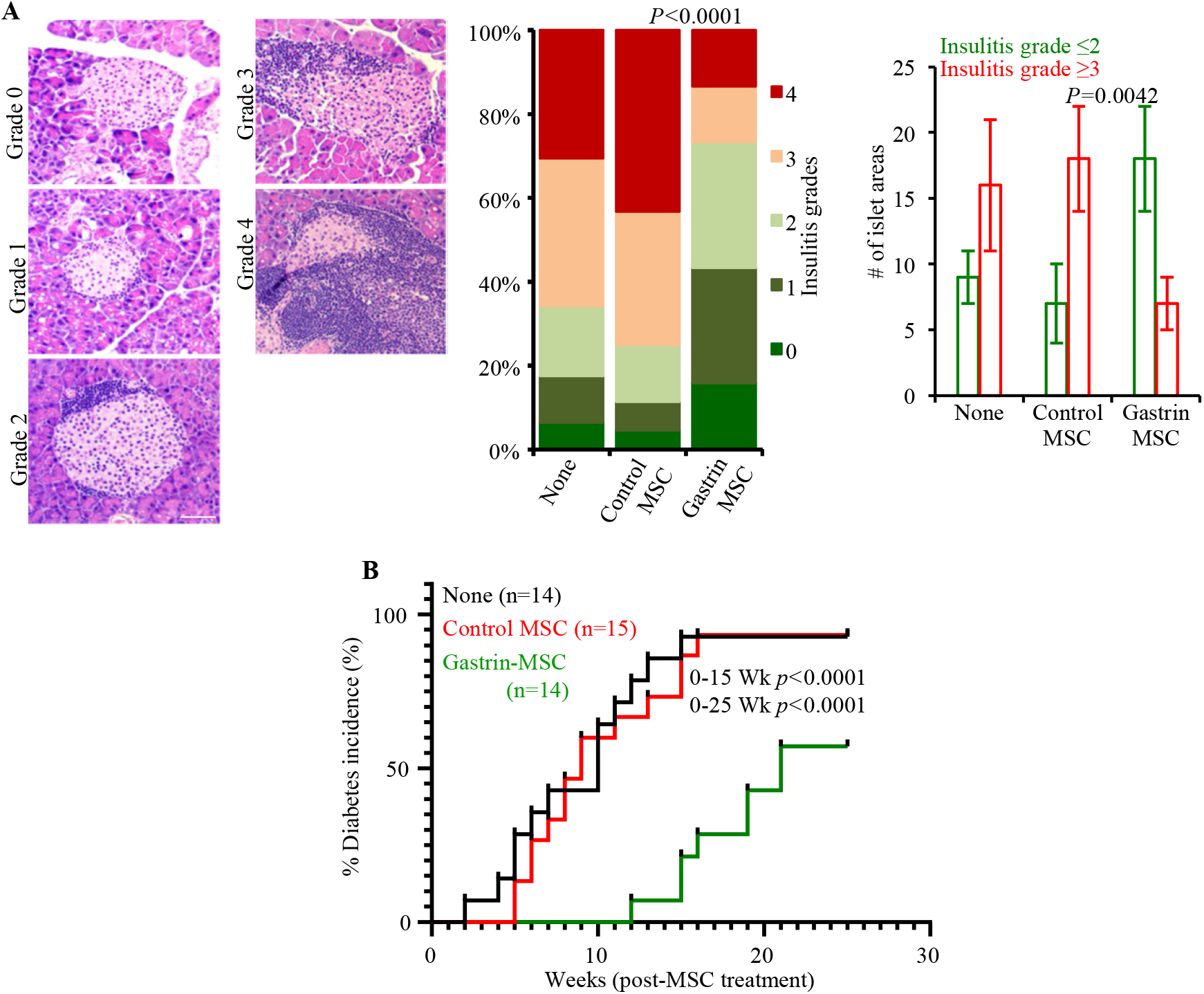
Impact of Gastrin MSC treatment on T1D incidence and insulitis. Ten-to 12-week-old pre-diabetic female NOD mice were left untreated (none) or injected with Control-MSCs or Gastrin-MSCs as described for Fig. 2. A) Cohorts of mice (a total of 6 mice/group) were euthanized 30 days post-treatment, pancreatic sections were subjected to H&E staining and grading for insulitis severity. Examples of islets with different insulitis grades (left panel) and the percentage medium and large islet areas with different insulitis grades (middle panel) are shown. Images were acquired using 20x objective. Scale bar: 50μm. Grades: 0 = no evidence of infiltration, 1 = peri-islet infiltration (<5%), 2= 5-25% islet infiltration, 3 = 25–50% islet infiltration, and 4 = >50% islet infiltration. A total of 150 areas (25 islet areas/pancreas) from 3 or more intermittent sections were examined for each group. Statistical significance was assessed by Fisher’s exact test comparing relative numbers of islets with insulitis grades ≤2 and ≥3 between groups. Average number of medium and large islet areas (with insulitis grades ≤2 and ≥3) per pancreatic tissue (3 intermittent sections/pancreas) are also plotted (right panel). **B)** Cohorts of mice were monitored for hyperglycemia by testing for blood glucose levels every week. Glucose: >250 mg/dl for two consecutive weeks was considered diabetes incidence, and cumulative results of multiple experiments, using 3-4 mice/group in each experiment (total of 14 to 15 mice/group), are shown. *p*-values of control-MSC vs Gastrin-MSC (0 to 15 week and 0 to 25 week) comparisons by log-rank test are shown.

### Gastrin-MSC treatment at pre-diabetic stage results in higher frequencies of insulin producing islets

To determine the effect of treatment using Gastrin-MSCs on the abundance of functional islets, insulin positive islet frequencies in mice that received MSC treatment at pre-diabetic stage were determined, 30 days after the treatment. Cryosections of pancreatic tissues of control-MSC and Gastrin-MSC treated mice were stained using anti-insulin antibody and the insulitis severity and total number of islet areas with and without insulin positive cells was assessed. Islet features were then graded depending on immune cell infiltration (based on DAPI) and insulin staining. As shown in **Fig. 4**., while majority of the islets (>90%), including those with profound immune cell infiltration, in Gastrin-MSC treated mice were insulin positive, >60% of pancreatic islets from control mice groups were insulin negative. Interestingly, insulin positive as well as insulitis free islet frequencies of Gastrin-MSC treated mice appear to be considerably higher (>20%) than that of control-MSC recipients, which is <5%. This result suggests that prevention of T1D in Gastrin-MSC treated mice is associated with protection/preservation of overall insulin producing islet mass and/or restoration of insulin expression in non-functional β-cells.

**Figure 4:**
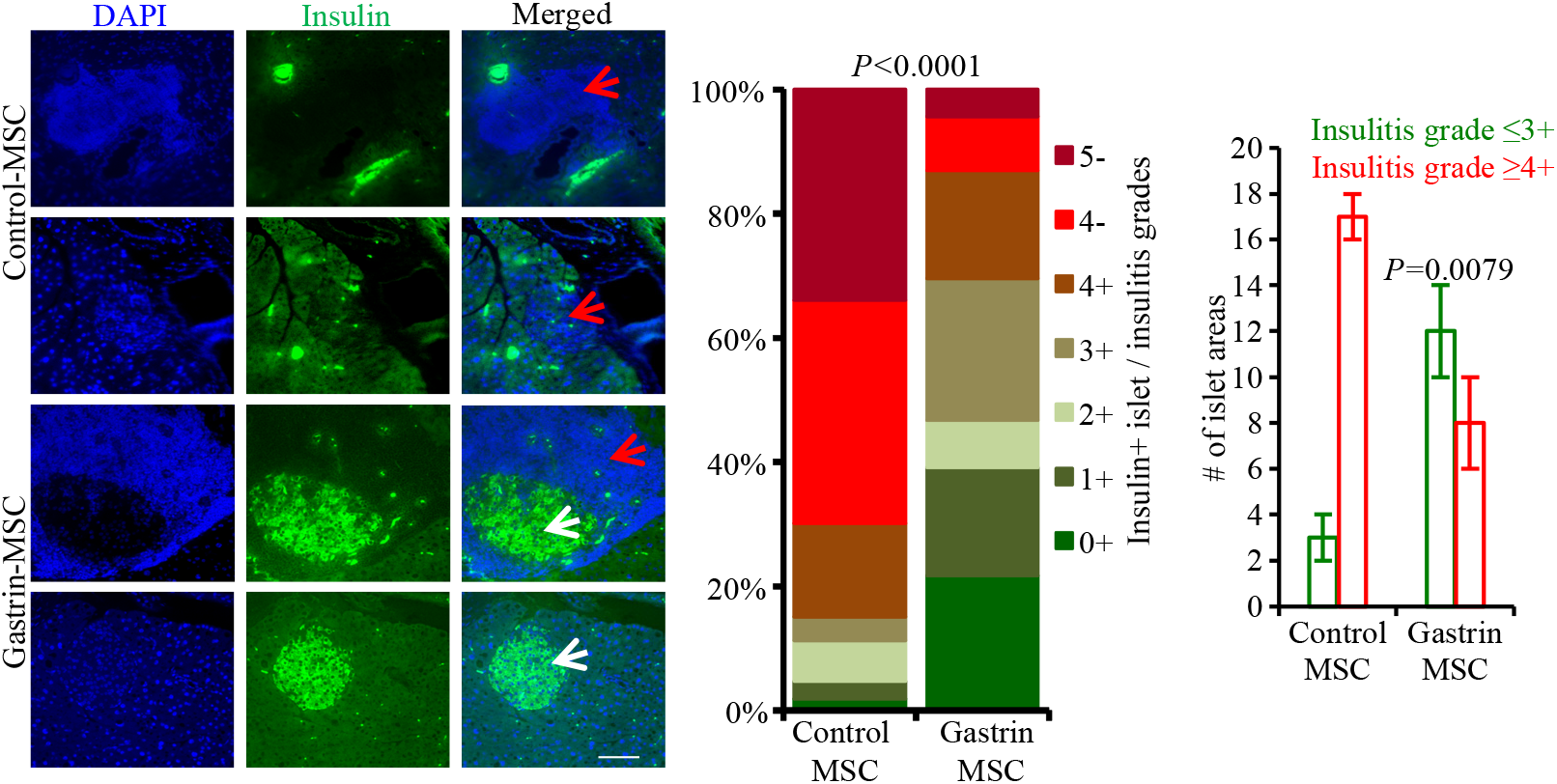
Impact of Gastrin-MSC treatment on insulin positive islet frequencies. Ten- to 12-week-old pre-diabetic female NOD mice treated with control-MSCs or Gastrin-MSCs as described for Fig. 2, euthanized after 30 days post-treatment, and pancreatic tissue sections were stained using anti-insulin antibody (green) and the nuclear stain DAPI (blue) and examined for insulitis and insulin positive islets. Representative microscopy images (acquired using 10x objective; scale bar: 100μm) indicating mostly insulin negative islet areas (red arrow) in control-MSC recipients and insulin positive islets (white arrow) in Gastrin-MSC treated mice are shown in the left panel. Islet features were graded 0-5 depending on immune cell infiltration (based on DAPI) and insulin staining (+ or −): 0+ (no infiltration/insulin+), 1+ (<5% infiltration/insulin+), 2+ (5-25% infiltration/insulin+), 3+ (25-50% infiltration/insulin+), 4+ (50-100% infiltration/insulin+), 4-(50-100% infiltration/insulin-), and 5-(only islet remnants left/insulin-), and percentage values of islets with specific grades among examined islet areas are shown in the middle panel. Medium and large islets, as well as islet/ductal remnant areas from multiple intermittent sections/pancreas (20 islet areas/pancreas) were examined for each group (total of 6 mice/group; 120 islet areas/group). *P*-value by Fisher’s exact test comparing relative numbers of islets with insulitis grades ≤3+ and ≥4+ between control-MSC recipients vs Gastrin-MSC recipients. Average number of medium and large islet/ductal areas (with insulitis grades ≤3+ and ≥4+) per pancreatic tissue are also plotted (right panel).

### Treatment of NOD mice with Gastrin-MSCs at early-hyperglycemic and diabetic stages results in transient reversal of hyperglycemia

Next, we examined if blood glucose levels and insulin producing islet abundance in mice with early-hyperglycemia and established diabetes can be altered by treating with Gastrin-MSCs. Female NOD mice with blood glucose between 140-250 mg/dl (early-hyperglycemic) and >250 (diabetic) mg/dl were injected once with control-MSCs or Gastrin-MSCs and monitored for blood glucose levels. As shown in **Fig. 5A**, while all control-MSC treated early-hyperglycemic mice (n=6) progressed to diabetic stage, all 4 early-hyperglycemic mice that received Gastrin-MSCs reverted to euglycemia (blood glucose <120 mg/dl) within 2 weeks and remained euglycemic for at least 6 weeks post-treatment. In the case of diabetic mice, Gastrin-MSC treatment caused transient reversal of hyperglycemia in mice that received the treatment when the blood glucose levels were at about 400 mg/dl or less, but not in those at overt-hyperglycemic stage. On the other hand, control-MSC recipient diabetic mice failed to revert to euglycemia. Of note, a separate experiment using only overt diabetic mice (glucose levels: >500 mg/dl; n=6 mice/group) showed no therapeutic effect either with control-MSCs or Gastrin-MSCs (not shown).

**Figure 5:**
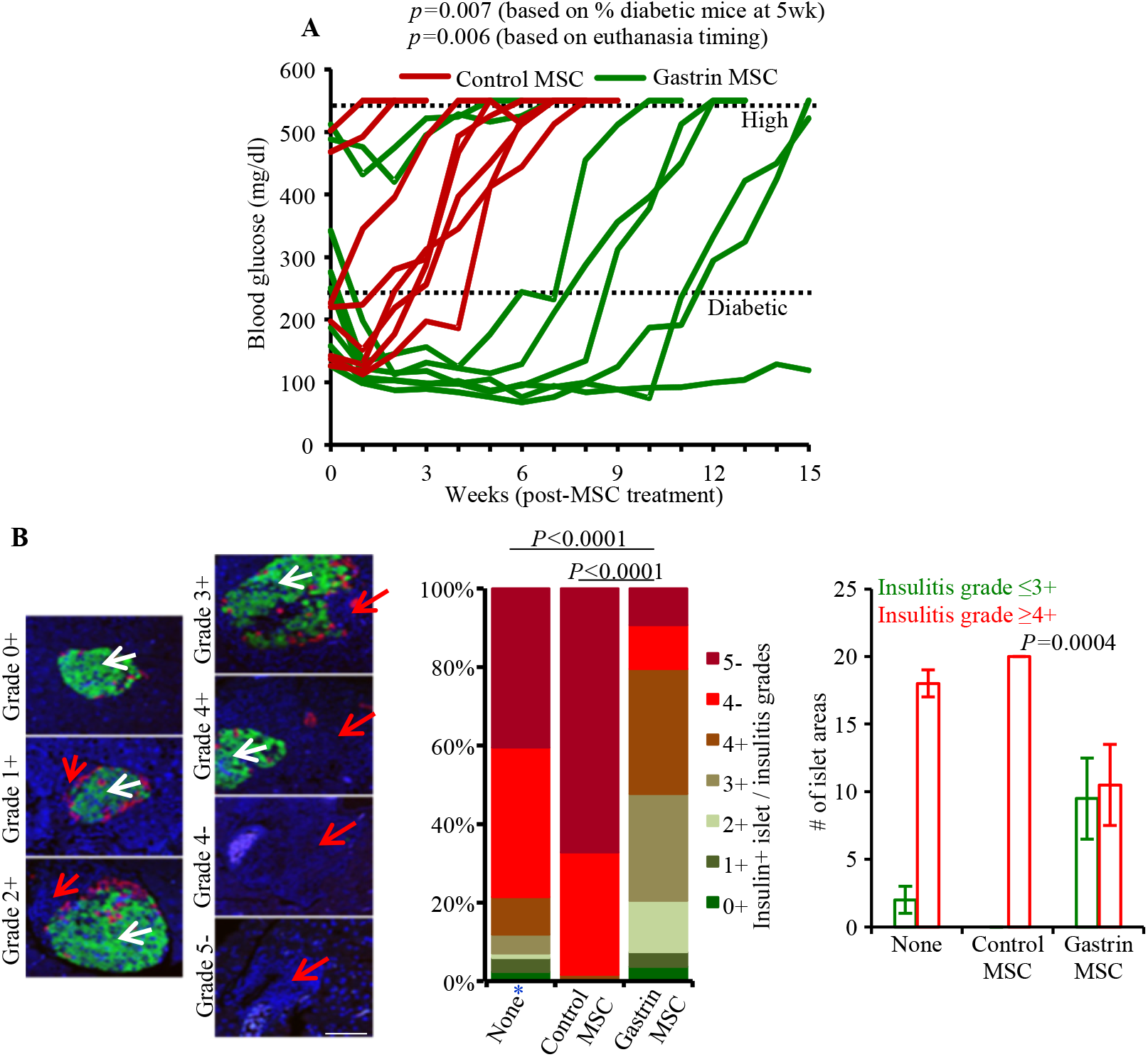
Impact of Gastrin-MSC treatment at hyperglycemic stages on blood glucose levels and insulin producing islet frequencies. Hyperglycemic group of female NOD mice (which included early-hyperglycemic and diabetic mice) were treated with control-MSCs or Gastrin-MSCs as described for Fig. 2. **A)** Blood glucose levels were examined before and every week after the treatment, and glucose values of individual mice are shown. Dotted lines represent “diabetic” and “high (endpoint)” glucose levels. *p*-values by Fisher’s exact test comparing the number of mice with diabetic glucose levels by week 5 post-treatment as well as by log-rank test comparing survival duration (euthanasia timing) of individual mice in each group. **B)** Cohorts of MSC treated early-hyperglycemic mice (5 mice/group) were euthanized 30 days post treatment, and pancreatic sections were subjected to insulin (green), glucagon (red) and DAPI (blue) staining and insulin positive islets and immune cell infiltration were scored as described for Fig. 4B. *Pancreatic tissues from a cohort of untreated mice that were at early-hyperglycemic stage at the time of euthanasia were included as “none” control group to assess the pre-treatment frequency of insulin positive islets. A total of 100 medium or large size islet/ductal areas with islet remnants (20 islet areas/pancreas) from 3 or more intermittent sections were examined for each group. *P*-value by Fisher’s exact test comparing relative numbers of islets with insulitis grades ≤3+ and ≥4+ between control-MSC recipients vs Gastrin-MSC recipients. Average number of medium and large islet/ductal areas (with insulitis grades ≤3+ and ≥4+) per pancreatic tissue are also plotted (right panel). Scale bar: 100 μm.

Pancreatic tissues were collected, at 30-days post-treatment, from cohorts of mice that were treated with control-MSCs and Gastrin-MSCs at early-hyperglycemic stage to examine insulitis and islet function. Pancreatic tissues from naïve, early-hyperglycemic (at the time of euthanasia) control mice were also collected and processed. Tissue sections were stained using anti-insulin and anti-glucagon antibodies and examined for the frequencies of “functional islets”. As shown in **Fig. 5B**, about 80% of islet areas, with or without immune cell infiltration, of Gastrin-MSC recipient mice showed some degree of islet function, as indicated by insulin and glucagon positivity, compared to about 2% insulin positive islet remnants in control-MSC treated mice. Of note, functional islet areas were detectable only in mice that received treatment at the early diabetic stages, but not at overt-hyperglycemic stage (not shown). These results show that reversal of hyperglycemia is possible by Gastrin-MSC treatment, if initiated soon after detection of clinical stage disease.

In a separate experiment, pancreatic tissues were harvested from a cohort of early-hyperglycemic mice 5 days post-eMSC injection and the sections were stained for cell proliferation marker Ki67 and insulin. **Supplemental Fig. 2** shows that albeit large number of islets with severe insulitis, significantly higher frequencies of insulin positive islets were detected in the pancreatic tissues of Gastrin-MSC treated mice, within 5 days of treatment compared to control-MSC recipients. This observation, although Ki67 staining did not produce conclusive results, suggests that higher frequencies of insulin positive β-cells in Gastrin-MSC treated mice could be, at least in part, the result of mitogenic activity of Gastrin, which involves the induction of insulin production in non-functional β-cells.

### Gastrin-MSC treated mice showed suppressed autoimmune response

While MSCs in general are thought to have immune modulatory properties[14–17], only the allogeneic or xenogeneic MSCs, but not the syngeneic MSCs showed immune modulatory properties and the ability to protect NOD mice from T1D upon repeated injections[14,15,34,37,51]. In fact, as shown in **Supplemental Fig. 3**, a single injection of syngeneic, non-engineered MSCs or control-MSCs had no impact on T1D progression in NOD mice. On the other hand, a single injection of non-engineered allogeneic MSCs caused a modest delay in T1D onset. Nevertheless, since T1D protection was achieved in NOD mice when treated with Gastrin-MSCs, we examined if the syngeneic control-MSC and Gastrin-MSC recipient mice showed differences in immune function. As observed in **Fig. 6A**, albeit modestly higher frequencies of IL10+ and IL17+ cells in Gastrin-MSC treated mice, differences in the frequencies of various T cell populations, including Foxp3+ cells, in the pancreatic LN of control- and Gastrin-MSC treated mice were not statistically significant. Similar observations were made when spleen cells were examined (not shown). Of note, control-MSC and Gastrin-MSC exposed, LPS-treated BM DCs showed comparable levels of activation markers such as CD80, CD86, CD40 and MHC II (**supplemental Fig. 4A**) suggesting that APC function of DCs is not impacted by Gastrin. Further, presence of Gastrin-MSCs in an antigen presentation assay did not have any significant impact on T cell proliferative, regulatory T cell or cytokine responses (**supplemental Fig. 4B & C**).

**FIGURE 6:**
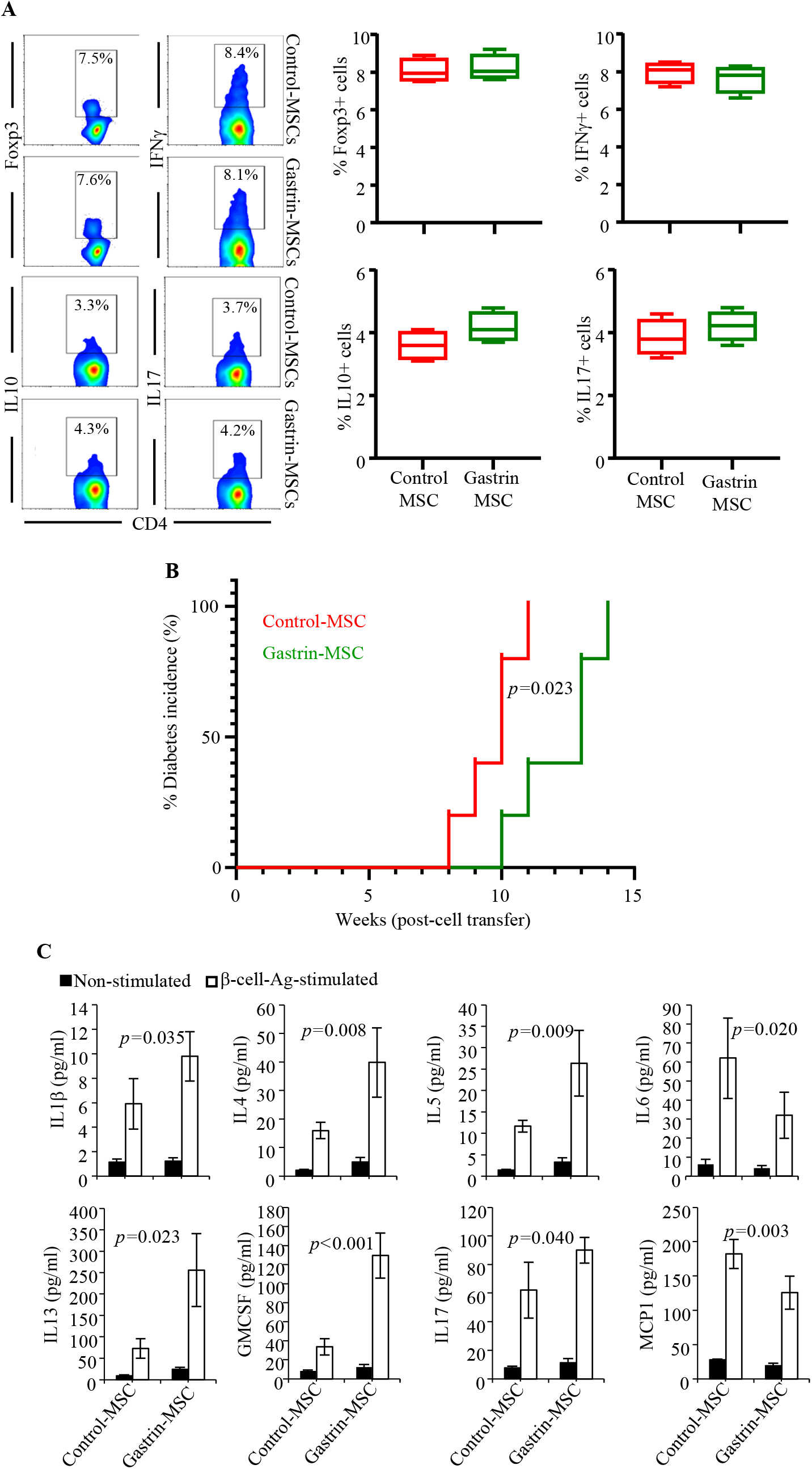
Impact of Gastrin-MSC treatment on immune cell phenotype and function. Prediabetic female NOD mice were treated with control-MSCs or Gastrin-MSCs as described for Fig. 2 and euthanized after 30 days to obtain PnLN cells. **A)** PnLN cells were stained for Foxp3, or for intracellular cytokines after brief stimulation using PMA and ionomycin in the presence of Brefeldin A, and analyzed by FACS. Representative FACS plots (left panel) and mean±SD values of 5 mice/group tested individually in duplicate (right panel) are shown. This experiment was repeated using similar number of mice. **B)** PnLN cells from at least 4 mice/group were pooled separately and injected i.v. into NOD-*Rag1-/-* mice (1×10^6^ cells/mouse; 5 recipients/group) and examined for blood glucose levels every week to detect hyperglycemia. *p*-value by log-rank test. **C)** PnLN cells (4 mice/group) from individual mice were cultured overnight in the absence or presence of β-cell Ag peptide cocktail for 48h and the supernatants were tested for cytokine levels by Luminex multiplex cytokine assay, and the mean±SD values of cytokine concentrations were plotted. *p*-value by Mann-Whitney test. Manufacturer suggested lower detection limits: IL1β, 3.2pg/ml; IL4, 6pg/ml; IL5, 2.1pg/ml; IL6, 10pg/ml; IL13, 3.8pg/ml; GMCSF, 13pg/ml; IL17, 8.7pg/ml; MCP1, 8.5pg/ml. This assay was repeated once using an additional 3 mice/group with comparable outcomes.

To determine the pathogenic effect of immune cells, PnLN cells from control- and Gastrin-MSC treated mice were activated using anti-CD3 antibody for 24h, and injected into NOD-*Rag1-/-* mice and monitored for hyperglycemia. As observed in **Fig. 6B**, recipients of PnLN cells from both control-MSC and Gastrin-MSC treated donors turned diabetic by 14 weeks of cell transfer. However, mice that received cells from Gastrin-MSC treated mice showed significantly delayed onset of hyperglycemia suggesting that the diabetogenic properties of immune cells are diminished in Gastrin-MSC treated mice. We, then, examined if self-antigen specific T cell responses are impacted by the eMSC treatment. PnLN cells from eMSC-treated mice were cultured ex vivo in the presence of β-cell antigen peptide cocktail[42–46], and the profiles of secreted cytokines were determined by employing Luminex multiplexed cytokine assay. **Fig. 6C** shows that cells from Gastrin-MSC treated mice, compared to that of control-MSC recipients, secreted significantly higher amounts of IL1β, IL4, IL5, IL13, IL17 and GM-CSF, and lower amounts of IL6 and MCP1 upon ex vivo activation using β-cell antigen. Interestingly, higher amounts of Th2 and Th17 response related cytokines (IL4, IL5, IL13 and IL17 particularly) were produced by immune cells from Gastrin-MSC recipients suggesting a skewed T cell response in these mice compared to control-MSC treated mice. Overall, these results show that while Gastrin-MSC treatment does not exert widespread immune modulation, self-antigen specific response and pathogenic properties of T cells in the pancreatic microenvironment are impacted as a consequence of this treatment and the associated establishment or maintenance of euglycemia, potentially contributing to protection from the disease.

## Discussion

Gastrin, which is produced primarily by the stomach and duodenum cells, binds to cholecystokinin B receptor and stimulates the release of gastric acid from parietal cells [52,53]. This peptide hormone can also cause increased division and differentiation of cells in the acid-secreting mucosa [53–55]. Of interest, gastrin is also expressed in the pancreas at the embryonic stage as well as in adult pancreas under stress including hyperglycemia in type 2 diabetes (T2D) and cancer[38,39,56,57]. Transient high expression of Gastrin and its receptors in fetal pancreas, in a period of pronounced islet neogenesis[58,59], suggests its involvement in increasing islet and β-cell mass. In fact, multiple reports have shown that Gastrin can increase the pancreatic β-cell mass^[60–69]^. Gastrin can also synergize with other agents such as GLP-1 and EGF and impact islet/ β-cell function and regulate blood glucose[61,65,69–72]. The β-cell neogenesis inducing potential of Gastrin was demonstrated using pancreas specific transgenic mice as well as by Gastrin therapy[61,70]. While the systemic use of Gastrin has been explored for enhancing β-cell mass [69,71,73], repeated high-dose systemic administration of this peptide hormone can be inefficient due to its short half-life. Further, its persistent high systemic levels can be unsafe due to global side effects including unregulated cell proliferation and malignancies. Hence, approaches for preferential delivery of Gastrin to, and its low-level continuous production in, the pancreatic microenvironment are needed to enhance β-cell mass safely and effectively. In this study, we demonstrate that the MSCs that are engineered to exogenously produce Gastrin could be such delivery vehicles for protecting β-cell function to prevent, and potentially reverse, hyperglycemia in T1D.

It has been shown that MSCs can ameliorate tissue damage and improve function after lung injury, kidney disease, diabetes, myocardial infarction, liver injury, and neurological disorders^8,[19,74,75]^. Natural ability of MSCs to migrate in the body has also been described^[23,24,76]^. Of particular interest, for delivering desired protein factors to pancreas, i.v. injection of MSCs results in their specific migration to a site of injury or inflammation^[23,25,77]^. Therefore, the inherent property of MSCs to migrate to the site of inflammation makes them the ideal cell population for delivering protein factors to sites of inflammation. Engineering the MSCs exogenously and stably to express the protein factors of interest is expected to ensure the prolonged delivery of factors such as Gastrin. Unlike other stem cells that are difficult to engineer with viral vectors without affecting their properties, MSCs can be readily transduced with most available viral vector systems to produce secreted, surface and cytoplasmic proteins of interest^[78,79]^. Importantly, MSCs have been engineered to either augment their own natural production of specific proteins or to enable them to express other therapeutic proteins for various clinical conditions^[80–82]^. Furthermore, eMSCs have been effectively used to enhance their ability to repair tissues including myocardium, CNS and liver in experimental models^[83–85]^. With respect to T1D therapy, eMSCs that express Pdx1 and other β-cell specific transcription factors have shown the ability to become insulin producing cells,in vitro and in vivo^[86–88]^. While, how effective these MSC preparations are in differentiating into IPCs in an autoimmune microenvironment and if they can reverse the established disease is largely unknown, these reports show the possibility that MSCs from T1D patients and pre-clinical models, despite their defective differentiation and immunomodulation properties[34,36], can be exploited as delivery vehicles for proteins of interest. Therefore, we postulated that if MSCs that are engineered to secrete Gastrin can, upon injection, reach and condition the pancreatic microenvironment, then these cells will be effective in increasing insulin producing cells and reversing hyperglycemia without being reliant on their multi-potent and immune modulatory properties. In fact, our observations that: 1) engineered-MSCs, upon inoculation, reach the pancreatic islets with insulitis, 2) Gastrin production can be detectable, albeit at low levels, in the pancreatic tissue of Gastrin-MSC recipient NOD mice even after 30 days of injection, 3) insulin expressing islet frequencies are higher, and 4) diabetes can be prevented and early hyperglycemia can be reversed for significant durations, do support this notion.

With respect to islet β-cell mass and insulin production in T1D, two key Gastrin associated effects have been reported. First, it has shown that Gastrin’s mitogenic property not only triggers increased insulin production by residual β-cells, but it alone or along with other factors such as GLP-1, EGF, and TGFα also induces the proliferation of these cells[38–41,61,65,69–72]. Second, studies using pancreas specific Gastrin and TGFα transgenic mice, as well as that involving pancreatic ductal ligation, have shown that Gastrin can contribute to trans-differentiation of the exocrine acinar cells to duct-like cells as well as neogenesis of insulin producing islets from ductal cells[62,64,68,70]. Therefore, while the mechanism of Gastrin-MSC treatment induced protection of NOD mice from T1D in our study is not known, it is possible that higher insulin positive islets found in these mice as compared to control MSC-recipients are, at least in part, due to the direct effects Gastrin on pancreatic tissues. Substantiating the likelihood of direct impact on pancreatic islets, we observed that many of the islet structures with severe insulitis in Gastrin-MSC treated mice, unlike in control MSC treated mice, were insulin positive as early as 5 days after the treatment. This suggests the possibility that Gastrin through its mitogenic activity induces insulin expression in non-functional, residual β-cells even in profoundly inflamed islets. We also observed that the insulin positive islet frequencies of Gastrin-MSC recipients, 30 days post-treatment, was significantly higher than that of a cohort of pre-treatment stage mice. With respect to the mechanism of Gastrin-MSC treatment associated increase in islet/β-cell function, it is possible that mitogenic activity of eMSC delivered Gastrin causes proliferation of residual β-cells, and β-cell neogenesis and/or trans-differentiation from other islet cells and ductal cells. These aspects need to be studied systematically in the future.

Notably, syngeneic Gastrin-MSC treated NOD mice showed suppression of autoimmune and pro-inflammatory response to a certain degree. While the previous report showing the need for repeated injections with syngeneic MSCs[34], and our observations that a single dose of syngeneic non-engineered or control-MSCs had no impact on T1D incidence in NOD mice, suggest that the immune modulation observed in Gastrin-MSC recipients is treatment associated. A direct effect for Gastrin on immune cells is largely unknown. Our observations that immune function of Gastrin-MSC treated mice, compared to control-MSC treated mice, is skewed towards Th17 and Th2 types suggest that Gastrin-MSC treatment does impact immune function, perhaps indirectly through causing changes in the pancreatic microenvironment. In this regard, our in vitro coculture assays using control and Gastrin-MSC and NOD spleen cells and purified immune cells showed comparable proliferative and cytokine responses. Furthermore, although it has been shown that Gastrin-receptor (cholecystokinin) is expressed on immune cells such as monocytes[89,90], we did not observe the expression of this receptor on immune cells (not shown), any significant impact of Gastrin on activation marker expression on DCs or on T cells during antigen presentation by DCs. It is possible that, as suggested in previous reports[91–94], hyperglycemia alone can impact various physiological functions including gut permeability and cause systemic pro-inflammatory response aggravating the disease complications. Hence, an explanation for the observed in vivo immune modulation in Gastrin-MSC recipients is that Gastrin-MSC treatment enhanced the β-cell function (potentially due to the mitogenic effect of Gastrin) and suppressed hyperglycemia, contributing to modulated immune cell function and the overall autoimmune and disease outcomes. In fact, our observations that Gastrin-MSC recipients show profoundly higher degree of insulin positivity even among insulitis positive islets within a short period as early as 5 days post-treatment supports this notion.

Overall, our study suggests that the use of autologous MSCs to deliver peptide hormone and β-cell mitogen, Gastrin in the pancreatic microenvironment can be an effective and safer approach for preventing T1D and reversing new onset hyperglycemia. It is possible that repeated injections as well as combination treatment approaches involving other islet regeneration and immune modulatory factors could potentially enhance the efficacy of this approach for achieving stable/permanent euglycemia, even in patients with clinical stage disease.

## Data availability

The datasets generated and/or analyzed during the current study are available from the corresponding author on reasonable request.

## Resource availability

The cDNA vectors generated during the current study is available from the corresponding author on reasonable request.

## Author contribution

M.G. researched and analyzed data and edited the manuscript, R.G. researched and analyzed data and edited the manuscript, G.L, researched data, B.J. researched data, and C.V. designed experiments, researched and analyzed data, and wrote/edited manuscript.

## Acknowledgements

This work was supported by unrestricted research funds from MUSC and National Institutes of Health (NIH) grants R21AI069848, R21AI133798 and R01AI073858 to C.V. C.V. is the guarantor of this work and, as such, has full access to all the data in the study and takes responsibility for the integrity of the data and accuracy of the data analysis. The authors are thankful to Cell and Molecular Imaging, Pathology, Proteomics, immune monitoring and discovery, and flow cytometry cores of MUSC and UIC for the histology service, microscopy, FACS and multiplex assay instrumentation support.

