## supplemental figures for "Gastrin producing syngeneic mesenchymal stem cells protect non-obese diabetic mice from type 1 diabetes"

Supplemental Fig. 1

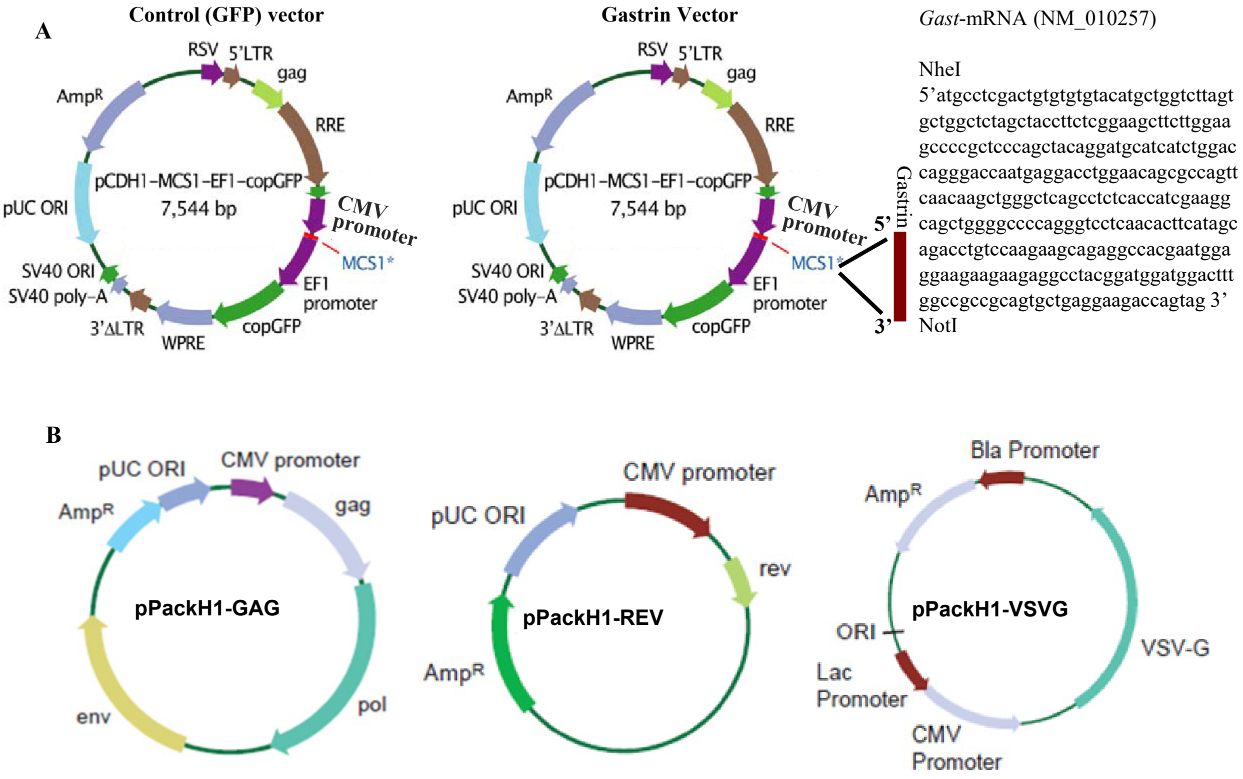


**Supplemental figure 1:** Lentiviral vector system that was used for engineering MSCs in this study.

Supplemental Fig. 2

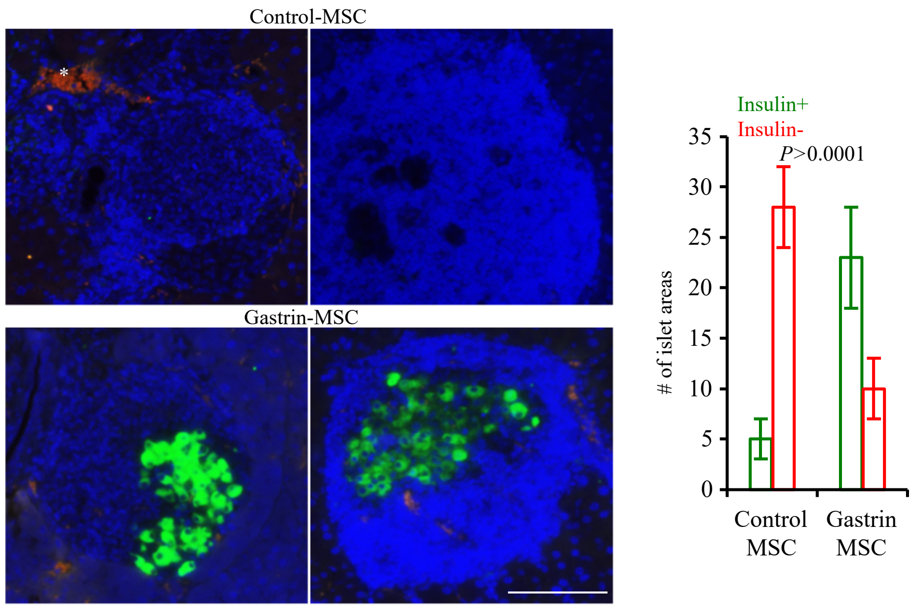


**Supplemental figure 2: Impact of Gastrin-MSC treatment of early-hyperglycemic mice on insulin positive islet frequencies.** Early-hyperglycemic female NOD mice were treated with control-MSCs or Gastrin-MSCs (3 mice/group) as described in Fig. 3 and the mice were euthanized on day 5, pancreatic tissues were embedded in paraffin and the sections were subjected staining using anti-insulin (green) antibody and DAPI (blue). Images of representative islet areas of each group are shown (left panel). Insulin positive and negative islet areas of multiple intermittent sections of each tissue (at least 30 islet areas/tissue) were counted and average values are shown in the right panel. Scale bar: 100 µm. While insulin expression was detected in most islet areas of mice that received Gastrin-MSCs, control-MSC recipient mice showed very few insulin positive islets areas, potentially due to the mitogenic activity of Gastrin in terms of triggering insulin expression in residual β cells.  This notion needs to be tested extensively in the future.

**Note:** These sections were also stained using anti-Ki67 antibody (red). However, red fluorescence in these images does not appear to be specific or overlap with DAPI staining, especially in the vascular area of upper left image(*). Hence, Ki67+ islet areas were not quantified. Future studies will address if Gastrin-MSC treatment induce islet cell proliferation.

Supplemental Fig. 3

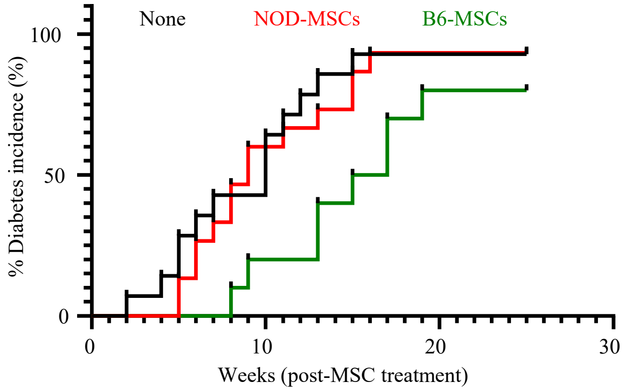


**Supplemental figure 3: Impact of treatment using syngeneic and allogeneic MSCs on T1D in NOD mice.** Ten- to 12-week-old pre-diabetic female NOD mice were left untreated (none) or injected with non-engineered syngeneic NOD mouse MSCs or allogeneic B6 mouse MSCs ($1x10^6$ cells/mouse) and monitored for hyperglycemia. n=10 mice/group.

Supplemental Fig. 4

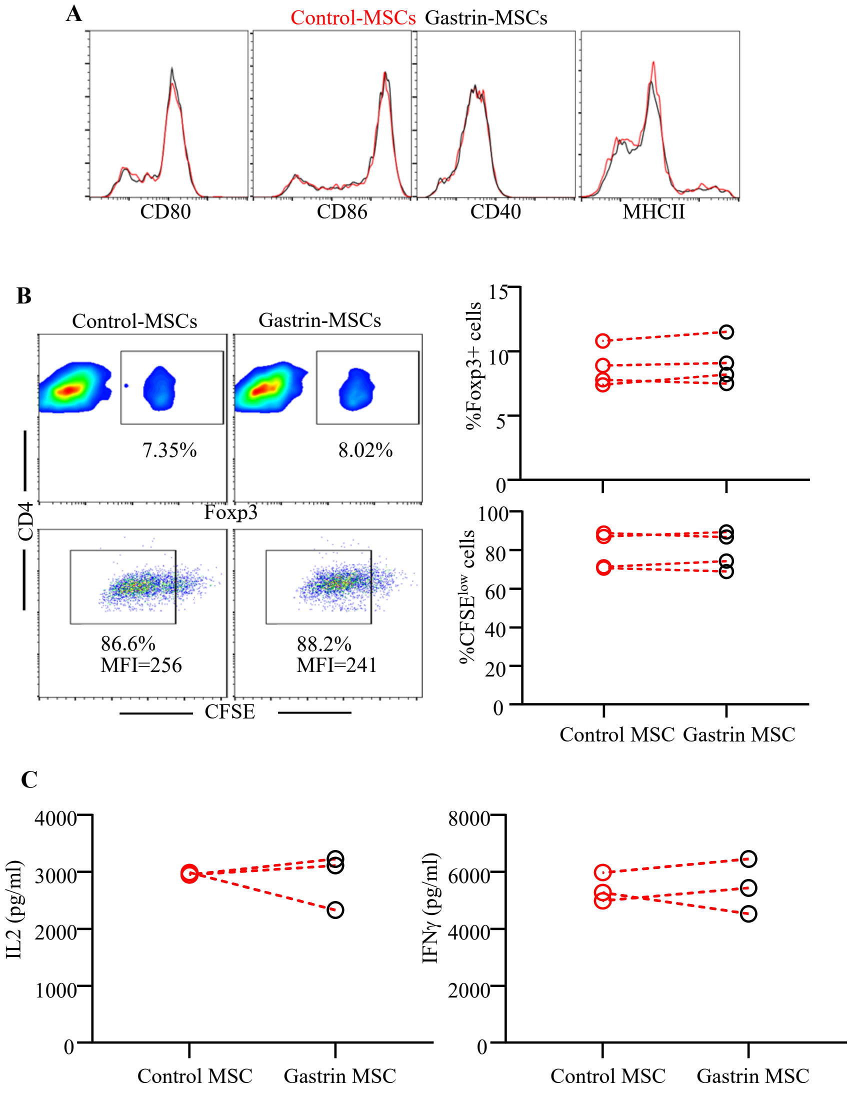


**Supplemental figure 4: Gastrin does not have significant influence on DC activation and antigen presentation.** BM DCs ($1x10^6$ cells/ml) were cultured in the presence of bacterial LPS (1 μg/ml) and control-MSCs or Gastrin-MSCs ($2x10^5$ cells/ml) for 48h. **A)** Cells were examined for activation marker levels by FACS. CD11c+ cells were gated for the overlay histograms. **B)** LPS activated DC and MSC mixture ($5x10^5$ cells/ml) were cultured in the presence of unlabeled or CFSE-labeled CD4+ T cells ($2x10^6$ cells/ml) from NOD-BDC2.5 TCR-transgenic mice for 4 days and examined for CFSE dilution and Foxp3+ T cell frequencies by FACS. CD4+ cells were gated for the analysis. **C)** supernatants from the cultures of panel B were tested for various cytokine levels by multiplex assay and the values of IL2 and IFNγ are shown. Representative flow graphs and/or values from three independent assays, each performed in triplicate, are shown.
